# sciCAN: Single-cell chromatin accessibility and gene expression data integration via Cycle-consistent Adversarial Network

**DOI:** 10.1101/2021.11.30.470677

**Authors:** Yang Xu, Edmon Begoli, Rachel Patton McCord

## Abstract

The booming single-cell technologies bring a surge of high dimensional data that come from different sources and represent cellular systems from different views. With advances in single-cell technologies, integrating single-cell data across modalities arises as a new computational challenge and gains more and more attention within the community. Here, we present a novel adversarial approach, sciCAN, to integrate single-cell chromatin accessibility and gene expression data in an unsupervised manner. We benchmarked sciCAN with 3 state-of-the-art (SOTA) methods in 5 scATAC-seq/scRNA-seq datasets, and we demonstrated that our method dealt with data integration with better balance of mutual transferring between modalities than the other 3 SOTA methods. We further applied sciCAN to 10X Multiome data and confirmed the integrated representation preserves information of the hematopoietic hierarchy. Finally, we investigated CRSIPR-perturbed single-cell K562 ATAC-seq and RNA-seq data to identify cells with related responses to different perturbations in these different modalities.

---

Within the last decade, single-cell technologies have advanced our understanding in a broad range of biological systems. Single-cell RNA-seq and single-cell ATAC-seq, along with other single-cell assays, have revealed distinct cellular heterogeneity at a comprehensive level, from genomic variations to epigenomic modifications and transcriptomic regulation^1-5^. Analyses based on single-cell data have also provided reliable databases for biomedical research and valuable references for medical discovery. As the number of single-cell omics datasets grows, there is increasing demand for fast and accurate computation. Consequently, deep learning has become a trending topic in single-cell data analysis. Much recent research has focused on developing reliable and fast deep learning tools to accommodate the scaling demand, such as cell-type annotation^6^, doublet identification^7^, data de-noising^8^, and batch correction^9^.

Among all applications of deep learning in single-cell analysis, data integration remains one of the grand and rising challenges in the community^10,11^. Many different single-cell RNA-seq platforms were simultaneously and rapidly developed, leading to an initial focus on methods to integrate datasets from different platforms. Batch effects are usually the most prominent variation when datasets from different sources are collected for integrative analysis but often are not biologically relevant. Single-cell databases confounded by batch effects are not applicable for general use. Therefore, removing batch effects is a critical step for revealing true biological variation and necessary for building batch-invariant and applicable databases. So far, multiple methods have been proposed to address this problem^9,12-16^. Among these integration methods, deep generative models were also extensively tested in single-cell analysis and demonstrated their efficacy of learning discriminative representation from the original high dimensional space. The most common generative models are Variational Autoencoder (VAE). Variants of VAE models, which differ in their sampling approaches, have been proposed to learn representations for single-cell gene expression data^9,17-20^. The core component of VAE is the use of reconstruction loss, which encodes a sample in a representation that is drawn from a certain distribution, for example, a Gaussian distribution. The use of reconstruction loss also has an advantage of mapping noisy data to high-quality data, which further extends the ability of generative model to de-noise data or impute gene expression. Instead of using VAE to learn representation for single-cell RNA-seq data, two research groups simultaneously modified VAE to address batch effects using an adversarial approach^19,20^. Two methods, named scGAN and AD-AE, respectively, used generative adversarial network (GAN) as the main framework for learning the latent space that is not entangled with batch effects. Starting from a VAE model, both scGAN^19^ and AD-AE^20^ introduced adversarial domain loss into the generative model and transferred the learning from reconstruction of data to diminishing of non-biological variation. This approach turned out to be effective in removing batch effects within single-cell gene expression data. Previous work has only focused on the use of adversarial learning in single-cell RNA-seq data.

Considering the success of deep generative models in batch-effect correction, we extended its use to single-cell data integration across different modalities. In this study, we focus on modality differences and developed an improved adversarial domain adaption approach to address multimodal data integration for chromatin accessibility (ATAC-seq) and gene expression (RNA-seq) data. Our method differs from both scGAN and AD-AE in that it uses a cycle-consistent adversarial network to learn the joint representation for both chromatin accessibility and gene expression data^21^. We term our method sciCAN (***s***ingle-***c***ell chromatin accessibility and gene expression data ***i***ntegration via ***C***ycle-consistent ***A***dversarial ***N***etwork), which removes modality differences while keeping true biological variation. We previously developed a deep learning method, SMILE, to perform integration of multimodal single-cell data^22^. SMILE requires cell anchors for integration. This limits the use of SMILE in cases where corresponding cells are known across modalities. Different from SMILE, sciCAN doesn’t require cell anchors and thus, it can be applied to most non-joint profiled single-cell data. We first benchmarked our method with 3 SOTA methods in 5 ATAC-seq/RNA-seq datasets, and we demonstrate that our method deals with data integration with a better ability to transfer cell type labels in both directions between modalities than the other 3 SOTA methods. To demonstrate the method’s utility in integrative analyses, we applied sciCAN to joint-profiled peripheral blood mononuclear cells (PBMC) data by 10X Multiome platform and we confirmed that the hematopoietic hierarchy is conserved at both chromatin accessibility and gene expression levels. Finally, we investigated CRSIPR-perturbed single-cell K562 ATAC-seq and RNA-seq data, and we identified that some cells in both modalities share common biological responses, even though the two modalities were profiled with different gene perturbations. Combining the results above, we expect our work will fill the gap to allow generative models to be used in integrative analysis of multimodal single-cell data.

## Results

### Overview of sciCAN and potential applications

We first show the model architecture of sciCAN, which contains two major components, representation learning and modality alignment (Fig. 1a). Encoder *E* serves as a feature extractor that projects both high dimensional chromatin accessibility and gene expression data into the joint low dimension space. For representation learning, we use noise contrastive estimation (NCE) as the single loss function to guide *E* to learn the discriminative representation that can preserve the intrinsic data structure for both modalities. For modality alignment, we use two separate discriminator networks for two distinct uses. The first discriminator network *D*_*rna*_ is attached to *E* and is trained with adversarial domain adaptation loss. *D*_*rna*_ aims to distinguish which source the latent space *z* extracted by *E* comes from, while *E* is pushed to learn the joint distribution so that *D*_*rna*_ is less able to distinguish the modality source of latent space *z*. The second discriminator network *D*_*atac*_ follows a generator network *G* that generates chromatin accessibility data based latent space *z* from gene expression data. Adversarial training here will push *G* to find a connection between chromatin accessibility and gene expression data. Since the generated chromatin accessibility data is based on the latent space *z* of real gene expression data, the new latent space *z’* of generated data should align with its corresponding *z* of real gene expression data. Therefore, we add cycle-consistent loss as demonstrated in cycleGAN method to facilitate finding the connection between two modalities^21^. In practice, we build *E* with fully connected layers, which are followed by a batch normalization layer with Rectified Linear Unit (ReLU) activation. *D*_*rna*_ takes the 128-dimension *z* as input and forwards it through a three-layer multi-layer perceptron (MLP) to produce 1-dimension sigmoid activated output that predicts if the input *z* comes from single-cell RNA-seq data. Differently, *D*_*atac*_ takes output from *G* and forwards the input through a three-layer MLP to produce 1-dimension sigmoid activated output that predicts if input is generated by *G. G* is a decoder structure, which has two-layer MLP to restore dimension-reduced *z* to the original dimension of input data. Instead of calculating NCE directly on *z*, we further reduced *z* to 32-dimension output with linear transformation and 25-dimension SoftMax activated output, through two separated one-layer MLPs. This practice is the same as a recent method we developed for representation learning of single-cell data^22^. Once model training is done, we use encoder *E* to project both modalities into the joint representation for downstream analyses (Fig. 1b).

**Fig. 1:**
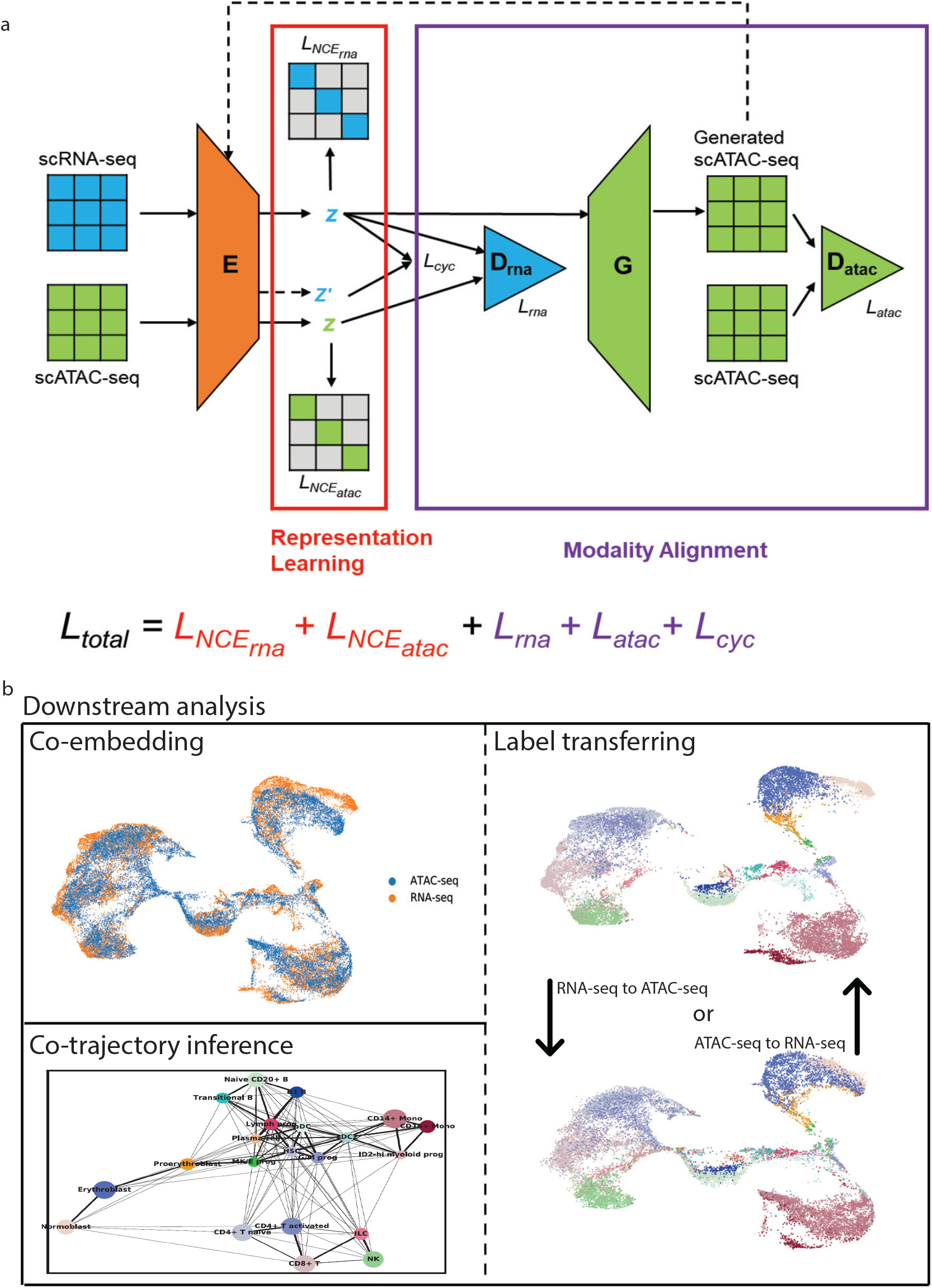
Overview of sciCAN and potential applications. a, sciCAN model architecture. sciCAN contains two major components, representation learning and modality alignment. The representation learning part of the model is highlighted in the red box, and the modality alignment part in the purple box. Inputs of scATAC-seq and scRNA-seq have been preprocessed to have the same feature dimensions, so they can share one single encoder *E*. The final total loss (*L*) is the sum of loss of representation learning in red and loss of modality alignment in purple. Of note, calculation of *NCE* is independent for scATAC-seq and scRNA-seq data. b, downstream integrative analyses can include but are not limited to co-embedding, co-trajectory, and label transferring.

### Benchmark of sciCAN with 3 SOTA integration methods

Our first metric (modality silhouette) evaluates how well two modalities align, and it directly reports whether discrepancy between chromatin accessibility and gene expression data is removed (maximum alignment gives a score of 0). In our benchmark test, we collected 5 datasets that come from distinct cellular systems. They are cell lines, human hematopoiesis, human lung, mouse skin, and mouse kidney, respectively. RNA-seq and ATAC-seq modalities may have different numbers of cells and even different numbers of cell types, except where both modalities were jointly profiled (Supplementary Table 1). Across 5 datasets, both sciCAN and Seurat integrated chromatin accessibility and gene expression data well, giving a smaller modal silhouette value. Following sciCAN and Seurat, LIGER also demonstrated a good overlapping between two modalities, but Harmony failed for the cell lines data (Fig. 2a). Though all 4 methods diminish the modality difference between chromatin accessibility and gene expression very well, it did not necessarily indicate that they learned the true structure that represents cellular components. Our second metric (cell-type silhouette) quantifies how well the joint representation reflects the data structure by distinguishing cell-types (in this case, a value of 1 is ideal). We used the author-reported labels as the ground truth. LIGER has the worst performance in human hematopoiesis and mouse skin data, Harmony is ranked as the worst in cell lines and human lung data, and Seurat scores the worst in mouse kidney data (Fig. 2a). Considering both good modality alignment and distinguishing cell-types, sciCAN shows the most consistently good performance across different experimental systems.

**Fig. 2:**
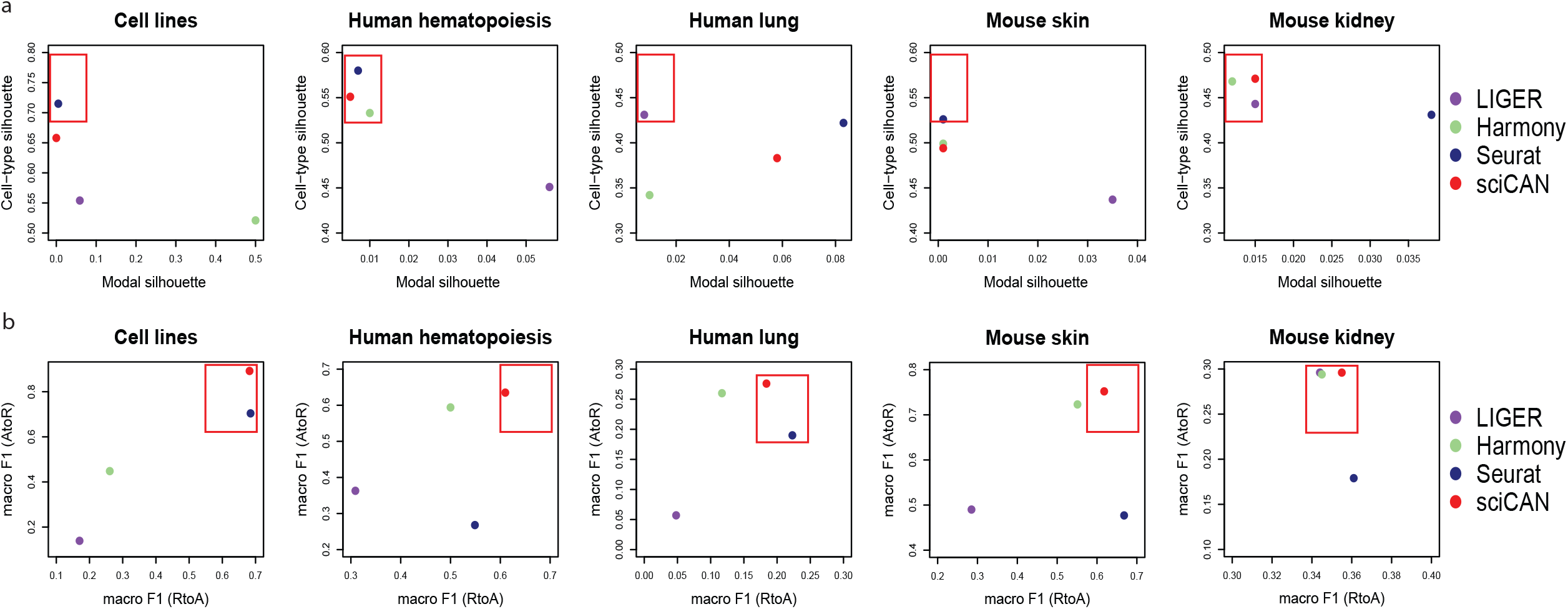
Benchmarking of sciCAN against 3 SOTA integration methods. a, Integration evaluation by modality and cell-type silhouette scores across 5 datasets. *x* axis corresponds to modality silhouette score while *y* axis to cell-type silhouette score. Ideal integration should be located in the top left corner of each dot plot. b, Integration evaluation by macro (upper panel) and weighted (lower panel) F1 scores across 5 datasets. *x* axis corresponds to label transferring from RNA-seq to ATAC-seq (RtoA) while *y* axis indicates label transferring from ATAC-seq to RNA-seq (AtoR). Ideal integration should be located in the top right corner of each dot plot.

Next, we focused on label transferring. Here, our goal is that the user could rely on the integrated space to predict cell-type labels for data from a single modality, given availability of cell-type labels from the other modality. We found Seurat has overall the best performance for label transferring from RNA-seq to ATAC-seq, while sciCAN has overall the best performance of label transferring from ATAC-seq to RNA-seq (Fig. 2b). When transferring labels back from ATAC-seq to RNA-seq, Seurat turns out to be the worst among the 4 methods. This may relate to the design of Seurat. Different from the other 3 methods, Seurat inherently uses gene expression data as reference data and projects chromatin accessibility data to the gene expression space. In cell line data, Harmony failed to transfer labels either from RNA-seq to ATAC-seq or ATAC-seq to RNA-seq. Among 4 methods, LIGER shows the worst performance with regard to label transferring (Fig. 2b). Visual inspection of the integration also indicates that both Seurat and sciCAN show overall better integration performance than LIGER and Harmony (Supplementary Fig. 1-5).

The default architecture of sciCAN shown in Fig. 1 has RNA-seq data playing the central role, primarily because RNA-seq data usually shows greater discriminative power than ATAC-seq in terms of cell-type identification^22-26^. Next, we examined if this setup is critical to good integration by sciCAN. We switched the roles of RNA-seq and ATAC-seq data in the model training. Indeed, the ATAC-centered sciCAN model is consistently less accurate than RNA-centered sciCAN, suggesting discriminative representation learning benefits from taking advantage of the cell-type discriminative power of RNA-seq (Supplementary Fig. 6). Combining the results above, we conclude that the RNA-centered sciCAN shows consistently good integration performance across different cellular systems.

### Integration learned by sciCAN preserves hematopoietic hierarchy

The hematopoietic hierarchy has been extensively studied through single-cell analysis. Independent studies using scRNA-seq or scATAC-seq also confirmed that the relationships between cell types in this hierarchy is observed at both chromatin accessibility and gene expression levels^27-31^. Thus, hematopoietic data can be a good example for us to verify whether the integration learned by sciCAN is biologically meaningful. Instead of using scRNA-seq and scATAC-seq data that were profiled separately, we utilized jointly-profiled human PBMC dataset obtained through the 10X Multiome platform, which enables us to evaluate integration with ground truth. Blinding ourselves to cell pairing information, our first task is co-embedding RNA-seq and ATAC-seq and performing co-trajectory analysis to evaluate whether the joint representation learned by sciCAN preserves the hematopoietic hierarchy at both chromatin accessibility and gene expression levels. Indeed, PAGA, a trajectory inference tool for single-cell data, constructed a hematopoietic stem cell (HSC)-centered trajectory with the 128-dimension joint representation learned by sciCAN^32^. We also confirmed that progenitor cells surround the HSPCs and branch towards their differentiated cells, and their lineage commitments at both chromatin accessibility (Fig. 3a) and gene expression levels can be explained by the same gene signatures (Fig. 3b and Supplementary Fig. 7). In our second task, we borrowed and transformed the concept of RNA velocity into activity-expression velocity. In the original RNA velocity concept, positive velocity suggests an increase of unspliced transcripts followed by up-regulation in spliced transcripts^33,34^. This idea was further extended to velocity analysis of nuclear mRNA vs cytoplasmic mRNA^35^, and of Tn5 vs TnH accessible chromatin regions^36^. Here, we reframed this analysis into activity-expression velocity by using the idea that an increase of gene activity (accessibility) would come first, followed by an increase in gene expression. Given the joint representation, we predicted gene expression based on gene activity. Then, we used the true activity matrix and the predicted expression matrix to compute the activity-expression velocity with scVelo^37^. We also performed the same analysis using activity matrix and true expression matrix. We found that velocity computed with the predicted expression data resembles the velocity computed with true expression data (Fig. 3c), and this resemblance is likely to due to accurate expression prediction (Fig. 3d and Supplementary Fig. 8). Consistent to co-trajectory analysis, velocity with predicted expression data revealed that MK/E progenitor cells move towards erythroblasts while G/M progenitor cells towards monocytes (Fig. 3d and Supplementary Fig. 8). Combining the results in two tasks above, we concluded that sciCAN preserves meaningful biological information within the learned joint representation.

**Fig. 3:**
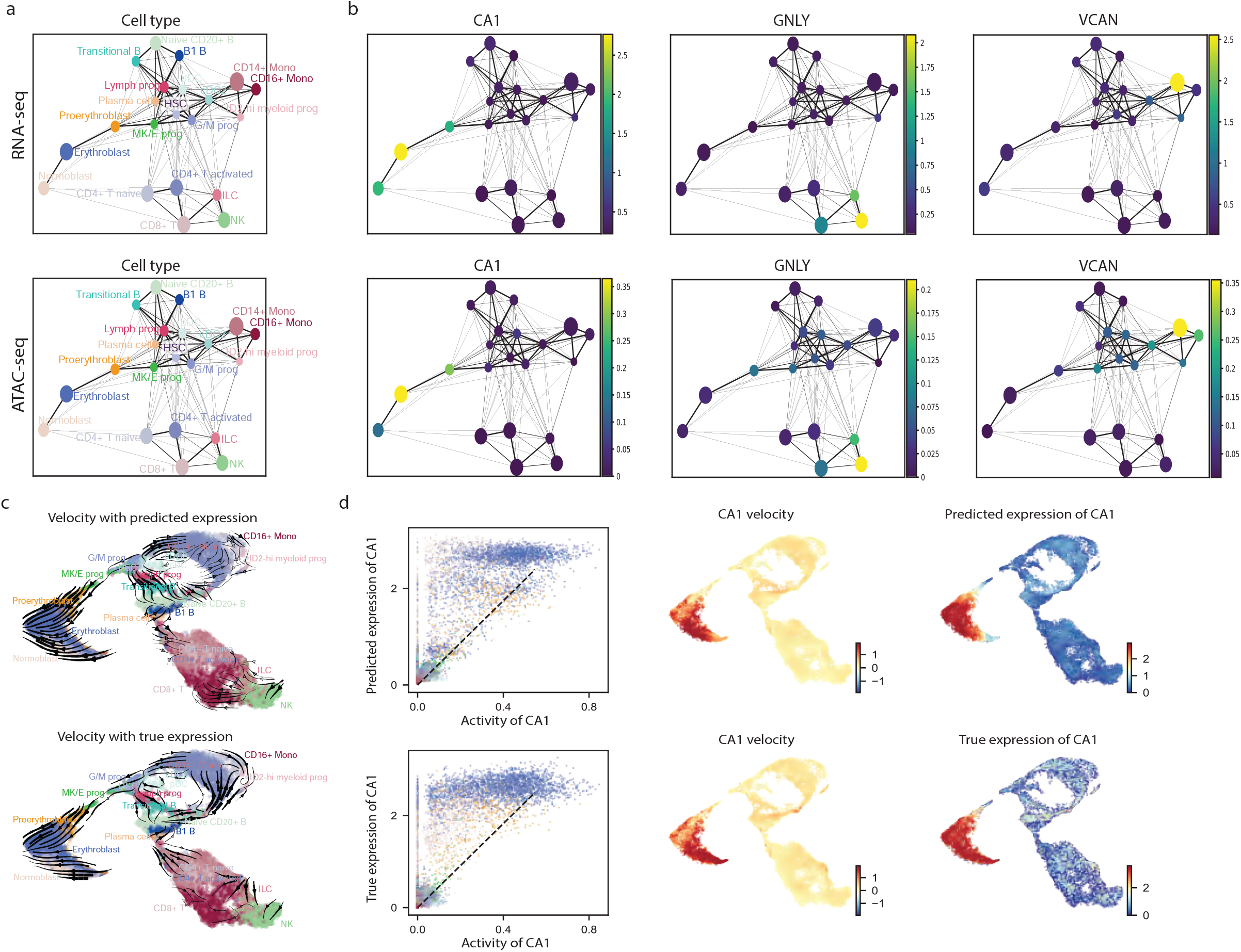
Integration learned by sciCAN preserves hematopoietic hierarchy. a, Co-trajectory analysis via PAGA using joint representation learned by sciCAN. Each dot is the sum of all cells annotated as the same cell type. Trajectory is visualized using RNA-seq (upper panel) and ATAC-seq (lower panel), separately. b, Enrichments of signature genes for 3 different lineages using both RNA-seq (top) and ATAC-seq (bottom) data. Colorbar indicates gene expression or gene activity level, respectively. c, Activity-expression velocity of the hematopoietic hierarchy. Velocity was calculated using predicted expression data (upper panel) or true expression data (lower panel). d, Activity-expression velocity of signature gene CA1 with either predicted expression data (upper panel) or true expression data (lower panel). Left: CA1 expression (predicted from ATAC-seq or measured by RNA-seq) is plotted vs. gene activity (accessibility) for each cell. Cell type indicated by color that corresponds to labels in previous panels. Dotted line indicates an estimated ‘steady-state’ ratio. Area above the dotted line suggests positive velocity, in which opening up of gene accessibility leads to up-regulation of its expression. Middle: the calculated velocity of CA1 superimposed onto the integrated representation across the hematopoietic hierarchy. Right: the expression of CA1 predicted by ATAC-seq vs. the true expression of CA1 superimposed onto the integrated representation.

### sciCAN identifies common responses after CRISPR perturbation

Combining single-cell sequencing with CRISPR enables a systematic examination of cellular response to genetic perturbation. Dixit et al. first introduced Perturb-seq to identify single-cell cellular response at the expression level after CRISPR perturbation^38^. Then, Perturb-ATAC was introduced to profile single-cell chromatin accessibility after CRISPR perturbation^39^. Nevertheless, a CRISPR-coupled joint-profiling single-cell assay has not been introduced. Therefore, multiple modality data integration is needed to determine how single cell responses to genetic perturbation compare at the transcriptomic and chromatin accessibility levels. As the final demonstration about potential application of sciCAN, we performed computational integration via sciCAN to create a joint view of cellular response after CRISPR perturbation. We selected single-cell K562 RNA-seq data by Perturb-seq and single-cell K562 ATAC-seq data through Spear-ATAC^38,40^. Notably these two studies used quite different sgRNA sets, sharing only 3 targets (*sgELF1, sgYY1*, and *sgGABPA*), so the integration cannot simply group like targets, but instead will be challenged to find similar biological responses to different gene perturbations. First, sciCAN enabled us to co-embed and co-cluster RNA-seq and ATAC-seq data, and we identified 3 distinct clusters (Fig. 4a). Next, we asked if the co-clustering makes sense in terms of gene signatures that lead to these clusters. Though the two studies used different sgRNA sets, we found gene activities of these 3 clusters have strong correlation to the gene expression profiles of the corresponding clusters in RNA-seq (Fig. 4b). Further, cells within each cluster shared gene signatures in both expression and accessibility (Fig. 4c). This suggests that cells may have similar response to different CRISPR-perturbations. Next, we ranked sgRNA targets for each cluster in both RNA-seq and ATAC-seq data. We found the 3 shared targets are in the top ranking in cluster 1 in RNA-seq but not ATAC-seq (Fig. 4d and Supplementary Table 1). We reason those cellular responses to perturbation at the chromatin accessibility level may be more variable than the responses at the gene expression level. Indeed, none of the ATAC-seq cell clusters have strongly dominant sgRNA targets as seen in the RNA-seq data. Therefore, we separated out cells that were targeted by the common targets *sgELF1, sgYY1*, and *sgGABPA* for a closer examination. We found that cells targeted by *sgELF1, sgYY1*, and *sgGABPA* in cluster 1 in both RNA-seq and ATAC-seq do have a distinct gene expression and activity signature compared to cluster 0 and 2, even though these cells were perturbed by the same sgRNAs (Fig. 4e). Shifting our focus to cluster 0 and 2, it is surprising that cells in these two clusters share the same top 5 sgRNAs (sgCEP55, sgOGG1, sgPTGER2 sgCAPBP7, sgCIT), in RNA-seq but are perturbed with completely different sgRNAs in ATAC-seq (Supplementary Table 2). To understand what makes cluster 0 and 2 different, we performed a differential gene activity test using cells targeted by the top 5 sgRNAs in cluster 0 and 2 ATAC-seq data. We then examined cells targeted by the shared top 5 sgRNAs in cluster 0 and 2 RNA-seq, and we found that the differential genes we identified through ATAC-seq could partially explain the different clustering of these cells in RNA-seq (Fig. 4f). Therefore, our integrated representation of these two independent datasets allows us to gain a better understanding of two subpopulations of cells that respond differently to the same gene perturbation.

**Fig. 4:**
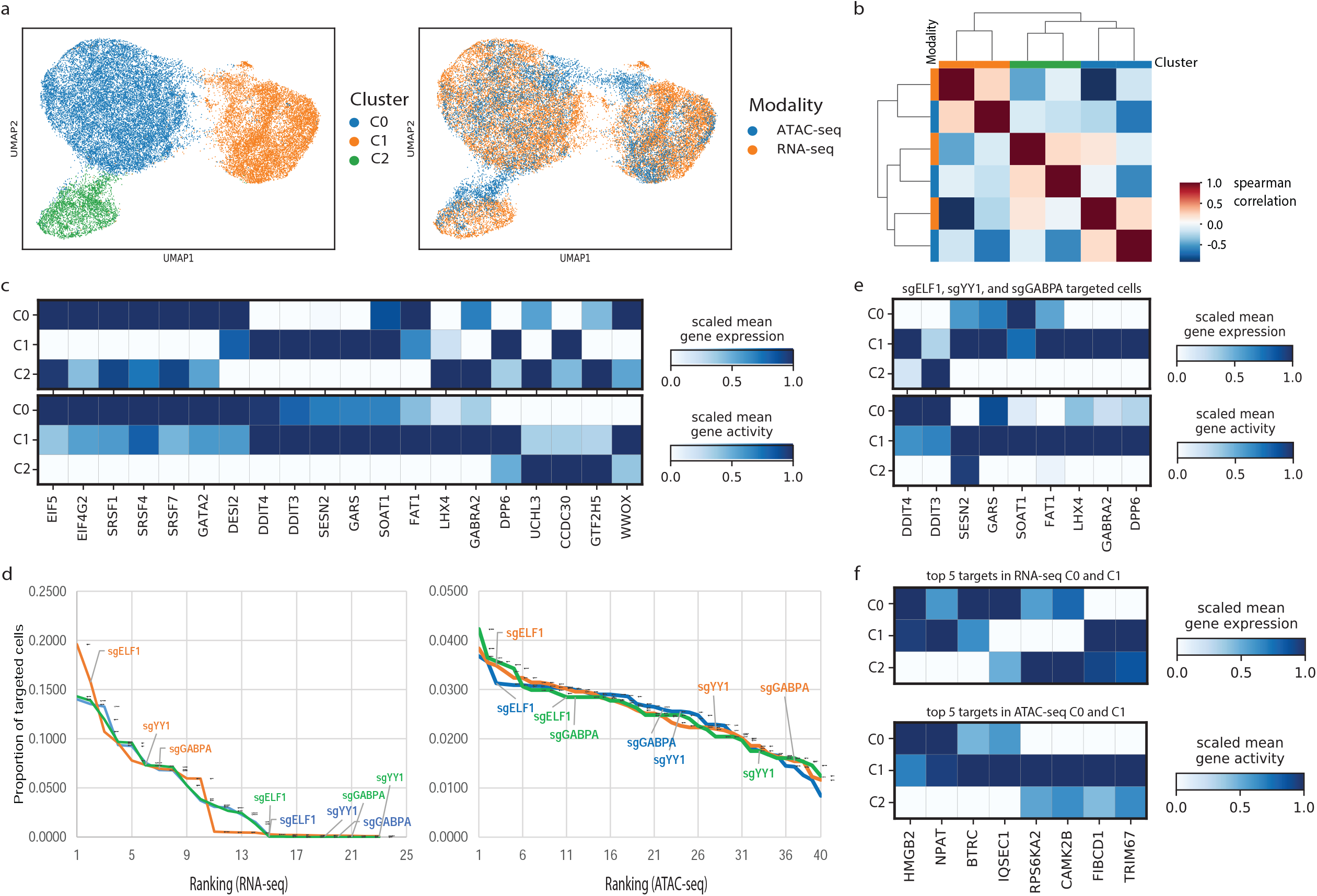
sciCAN identifies common response after CRISPR perturbation. a, Visualization of single-cell CRISPR-perturbed K562 RNA-seq and ATAC-seq data via UMAP. Cells are colored by identified cell clusters (left) and modality source (right). b, Spearman correlation between RNA-seq and ATAC-seq profiles of cells in different clusters in both modalities. Gene expression or gene activity matrix was averaged by cell clusters. c, Shared gene signatures of the 3 cell clusters in both modalities. Differential gene activities or expression were identified through ‘wilxocon’ test in Scanpy package. d, Ranking of sgRNA representation in each cluster (blue = C0, orange = C1, green = C2) in both RNA-seq (left) and ATAC-seq (right) data. Genes perturbed in both experiments are highlighted. e, Gene signatures of cells targeted by *sgELE1, sgYY1*, and *sgGABPA* in cell cluster 1. f, Genes whose activity patterns distinguish cells in cluster 0 and cluster 2 among cells in these clusters perturbed by the same gRNAs.

## Conclusion

In this study, we designed a novel adversarial approach for integration of single-cell chromatin accessibility and gene expression data. By benchmarking our method against the other 3 SOTA integration methods in 5 ATAC-seq/RNA-seq datasets, our showed that sciCAN and Seurat have overall superior performance of data integration. However, sciCAN shows good mutual label transferring either from RNA-seq to ATAC-seq or from ATAC-seq to RNA-seq, while this mutual information is lost via Seurat integration. In cases where researchers may want to translate ATAC-seq to RNA-seq for inferring gene expression, sciCAN would have an advantage over Seurat. We further demonstrated that sciCAN can be applied to different integrative analyses, like co-trajectory, activity-expression velocity, and co-clustering. All these results above demonstrate that sciCAN could empower integrative single-cell analysis for novel biological discoveries.

## Methods

### Representation learning

Deep metric learning has shown effective representation learning without supervision. Chen et al. used a simple framework to learn visual representations in a self-supervised manner^41^. They duplicated each image into two counterparts through image perturbation. The goal of learning is to maximize the consistency of any paired replicates in the latent space z. To achieve this goal, NCE is applied as loss function as shown in (1). In an N-sample batch, there will be *2N* samples through data augmentation, and each augmented image *i* has its corresponding counterpart *j* which is the same, despite the added image perturbation. Then, *cos* quantifies the cosine similarity of image *i* and *j/k* in the latent space *z*. Chen et al. demonstrated that this simple framework turns out to be a highly effective way to learn the discriminative representation without supervision. We adapted this approach in our previous study and showed the sample framework can produce discriminative representations for single-cell data^22^. Because of the property of this metric learning, our method is fully unsupervised. Users do not need to provide cell-type labels to start model training.

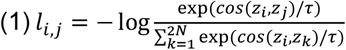

### Domain adaptation

Generative models with adversarial domain adaptation were successfully shown to transfer targets to source style and have general applications in image translation^42^. Recently, both scGAN^19^ and AD-AE^20^ incorporated adversarial domain adaption into a generative model for removing batch effects within single-cell expression data. For both studies, the goal is to find a batch-invariant representation for single-cell gene expression data from various sources. To achieve this, they stacked a discriminator to the encoder and trained the discriminator to distinguish which source the cell comes from using the latent space *z* projected by the encoder. Adversarial training, in this case, will push the encoder to approximate the joint distribution and become capable of projecting cells with data from different modalities to the same integrated representation. Here, we also used domain adaptation to train a discriminator to identify the modality source while the encoder is pushed to diminish modality difference.

### Cycle-consistent adversarial network

Besides the use of adversarial domain adaptation above, we further introduced a cycle-consistent adversarial part. This practice stems from a method called cycleGAN, which presented a SOTA outcome for the task of transferring image styles from one domain to another^21^. The success of establishing a connection between two image domains relies on the concept called “cycle consistency”. Starting from the original image, a generator network translates the image to the other domain. Then, a second generator network translates the image back to its original domain. Through this cycle, the translated-back image should be the same as the original image. Based on this information, adversarial training of generators can establish a reversible connection between two image domains. Different from the goal of cycleGAN, we aim to learn joint representation instead of translating chromatin accessibility to gene expression or vice versa. However, the fundamental concept is the same: we establish a cycle from encoder to generator, and from generator back to encoder. Then, the cycle-consistency loss is applied at the level of latent space *z*.

### Data preprocessing

All methods benchmarked in our study require anchoring genes for integration. We used a common practice that transforms the sparse ATAC-seq peak matrix to a gene activity matrix^23,43-45^. Here, we briefly explain the rationale behind this transformation. RNA-seq measures gene expression, so in a matrix of single-cell gene expression data, each row represents one cell, and each column contains expression values of one gene. The whole matrix represents gene expression levels of all genes across all cells. ATAC-seq, on the other hand, quantifies how accessible genomic loci are to regulatory proteins. Therefore, in a matrix of single-cell chromatin accessibility data, each row is one cell (the same as single-cell gene expression data) and each column contains accessibility values of one genomic locus. The sum of accessibility values of all genomic loci upstream of and within one gene body may relate to the potential of transcription of that gene. Therefore, to convert ATAC-seq data to a form that can be compared to RNA-seq data (a matrix of cells by genes), all accessibility peaks upstream of and within each gene body are summed to represent gene activity. In the converted gene activity matrix, each row is one cell, and each column is accessibility values of one gene. Therefore, after conversion, we can do a simple filtering and reordering to match features of chromatin accessibility and gene expression data. The Signac package provides this conversion process, and we ran the code available at https://satijalab.org/signac/articles/pbmc_vignette.html^45^. After we have both a gene activity matrix and a gene expression matrix, we normalize both modality data with (Log+1)-transformation, which adds 1 as a pseudo count to the matrix before log-transformation. Then, we identify the top 3000 highly variable genes (HVG) for each modality and use all identified HVG as features for integration. To identify the top 3000 HVG, we use Scanpy by calling the highly_variable_genes function^46^.

### Model training

We trained sciCAN in all datasets for 100 epochs. The learning rate starts from 0.005 with 0.0005 weight decay. All weights in the sciCAN model are updated through stochastic gradient descending. In the *NCE* loss function, temperature τ is a crucial parameter that affects discriminative power of the final representation. We set as τ = 0.15 for the 32-dimension linear-transformed output and τ = 0.5 for the 25-dimension SoftMax activated output, which is consistent to the practice in our previous study^22^. Detailed training code is also provided on sciCAN GitHub (https://github.com/rpmccordlab/sciCAN).

### Integration via LIGER

Multimodal single-cell data integration by LIGER was demonstrated in its published tutorial^14^. We used default parameters to perform integration of chromatin accessibility and gene expression data, and the final dimension of integrated representation by LIGER is 20 for all 5 benchmark datasets. Briefly, LIGER uses integrative nonnegative matrix factorization (iNMF) to identify metagenes that are shared between ATAC-seq and RNA-seq^47^. These metagenes are a weighted matrix of factor loadings of observed gene expression/activity. Then, cell loadings of these metagenes are used to perform joint clustering and other downstream analysis. Ideally, representations of cells from both modalities after iNMF should have been integrated in the same latent space and can be visualized via tSNE or UMAP^48,49^.

### Integration via Harmony

Harmony is the second integration method benchmarked in our study. Originally, Harmony was designed to correct batch effects within single-cell RNA-seq datasets^15^. Later, the novel use of Harmony in multimodal single-cell data integration was discussed in reviews^50,51^. Meanwhile, a batch-correction benchmark study showed that Harmony was ranked among the top 3 methods, with LIGER and Seurat, for integrating single-cell RNA-seq data^52^. Therefore, we included Harmony in our benchmarking of multimodal single-cell data integration. Harmony learns the joint representation through an iterative k-means clustering, and the outcome is a linear correction function that transforms the original principal components (PCs) to the batch-corrected PCs. Batch information is necessary to guide Harmony to distinguish what variation should be diminished during the k-means iterations. Principally, to integrate chromatin accessibility and gene expression data, modality information serves as the same role of batch information. Again, we used the default procedure of Harmony, in which we reduced the whole dataset into the first 30 PCs.

### Integration via Seurat

Seurat uses canonical correlation analysis to learn the shared latent space between two modalities. This approach is different from LIGER, Harmony, and our method, in a way that Seurat will first identify confident cell pairs between the two modalities. Then, Seurat uses these paired cells as anchors to learn a mutual neighborhood graph. Finally, it computes a projection that brings all other cells to this shared latent space. Because of its “anchor” design, Seurat needs pairwise computation of anchor points when datasets come from more than two sources. Since we only deal with the modality difference between chromatin accessibility and gene expression in this study, we do not need to perform pairwise computation of anchor points with Seurat. For benchmarking, we ran Seurat v3 with the tutorial on https://satijalab.org/seurat/archive/v3.0/atacseq_integration_vignette.html, and the final dimension of integrated representation by Seurat would be 50.

### Activity-expression velocity

Activity-expression velocity was calculated with scVelo^37^. We replaced the spliced layer with the gene activity matrix and the unspliced layer with the gene expression matrix, given the concept that increase of gene expression would follow increase in gene activity. To estimate first and second moments, we used the 128-dimension joint space learned by sciCAN, instead of PCA space.

### Data description

We collected unpaired chromatin accessibility and gene expression from 5 studies for benchmarking^53-57^. ATAC-seq and RNA-seq data in data 1 (Cell lines) and data 4 (Mouse skin) have the same number of cells and cell-types. This is because SNARE-seq and SHARE-seq simultaneously profile chromatin accessibility and gene expression features from the same cells^53,56^. Experimentally, each cell in ATAC-seq has its corresponding cell in RNA-seq in SNARE-seq and SHARE-seq data. However, in our study, we blind ourselves to this paired information for all 4 methods. For the other 3 datasets, data from ATAC-seq and RNA-seq were collected separately on different groups of cells. Therefore, they do not necessarily consist of the same number of cells and do not necessarily share the same cellular components. Cell-type annotations were also annotated separately by the authors, and the author-reported annotations serve as ground truth for integration evaluation. We also collected a joint-profiled human PBMC data by 10X Multiome platform to demonstrate that integration by sciCAN preserves biological information. Finally, we collected two independent CRISPR-perturbed single-cell K562 datasets that profiled chromatin accessibility^40^ and gene expression^38^, respectively. The brief description and citations for these 7 datasets are shown in Supplementary Table 1.

### Evaluation

To evaluate integration by each method, we proposed 4 metrics:

#### Modal and cell-type silhouette score

As we mentioned before, sciCAN reduces each dataset into 128-dimension spaces, while LIGER reduces the data to 20 dimensions, Harmony to 30, and Seurat to 50. Since final dimensions of the integrated representations by the 4 methods are not the same, we further used Uniform Manifold Approximation and Projection (UMAP) to reduce them into 2-dimensions with the same UMAP running parameters^58^. Then, we calculated modal and cell-type silhouette scores on the 2D UMAP spaces. A typical silhouette score *S* ranges from -1 to 1. To better reflect the integration outcome, we define modal silhouette as *abs(S)* and cell-type silhouette as (1 + *S*)/2. Of note, we used different labels to calculate modal and cell-type silhouette. For modal silhouette, the label used is modality information. A good integration should have chromatin accessibility and gene expression data largely overlapped. Therefore, 0 is the best outcome, and we ignore the positive/negative sign by using the absolute value of the typical silhouette score *S*. For cell-type silhouette, we used the author-reported annotation label to calculate *S* and then scale the output to the range from 0 to 1. Thus, cell-type silhouette 1 indicates the best integration that preserves cell-type structure.

#### F1 score from RNA-seq to ATAC-seq, and from ATAC-seq to RNA-seq

A useful integration of modalities should have the ability to transfer cell type labels from one datatype to another, either from RNA-seq to ATAC-seq or from ATAC-seq to RNA-seq. Given cell-type label availability from a single modality, the user should be able to predict cell-types for the other modality, with a fair accuracy. To evaluate how friendly the joint representation is for label transferring, we trained a Support Vector Machine (SVM) classifier with one modality and tested it with the other modality. The choice of SVM is simply based on a constant superior performance of SVM classifier across datasets. Then, we used macro F1 score to evaluate SVM classifiers trained with different joint representations by these 4 methods. Macro F1 score is the average of F1 scores for all cell-types, and it can help us reveal if integration is good for non-major cell-types. This is because cell-types are not balanced in most single-cell data and revealing non-major cell-types is critical for most single-cell analysis. A high macro F1 score can suggest that integration is also good for non-major cell-types.

## Data and code availability

All datasets used in our study are from previously published studies. The data accession in Gene Expression Omnibus or processed data link can be found in Supplementary Table 1. sciCAN code is provide on GitHub, with its own repository (https://github.com/rpmccordlab/sciCAN).

## Acknowledgements

This work was supported by NIH NIGMS grant R35GM133557 to R.P.M. We would also like to thank Heng Li for manuscript proofing and editing.

## Figure Legends

Supplementary Fig. 1: **Visualization of integration of Cell lines data**.

Visualization of integrated cell lines data via UMAP. Cells are colored by author-reported cell types (upper) and modality source (lower).

Supplementary Fig. 2: **Visualization of integration of Human hematopoiesis data**.

Visualization of integrated human hematopoiesis data via UMAP. Cells are colored by author-reported cell types (upper) and modality source (lower).

Supplementary Fig. 3: **Visualization of integration of Human lung data**.

Visualization of integrated human lung data via UMAP. Cells are colored by author-reported cell types (upper) and modality source (lower).

Supplementary Fig. 4: **Visualization of integration of Mouse skin data**.

Visualization of integrated mouse skin data via UMAP. Cells are colored by author-reported cell types (upper) and modality source (lower).

Supplementary Fig. 5: **Visualization of integration of Mouse kidney data**.

Visualization of integrated mouse kidney data via UMAP. Cells are colored by author-reported cell types (upper) and modality source (lower).

Supplementary Fig. 6: **Comparison of RNA-centered and ATAC-centered integration by sciCAN**.

Performances of RNA-centered and ATAC-centered sciCAN were evaluated by modality- and cell-type silhouette scores, and RtoA and AtoR macro F1 scores.

Supplementary Fig. 7: **Visualization of joint trajectory of hematopoiesis constructed through PAGA**.

Co-trajectory visualization without aggregating cells to one cell-type dot.

Supplementary Fig. 8: **Activity-expression velocity with predicted or true expression data**.

Activity-expression velocities of GNLY and VCAN. Velocities were calculated with both predicted and true expression data.

Supplementary Table 1: **Data description**.

Data used in this study are public. The number of available cells and annotated cell types are briefly described. The corresponding GEO accessions and links of processed data are also provided in this table.

Supplementary Table 2: **Ranking of sgRNAs in each cluster in both RNA-seq and ATAC-seq data**.

sgRNAs were ordered from the most to the least in each identified cluster for both RNA-seq and ATAC-seq.

