## Supplementary Figures and Tables for "sciCAN: Single-cell chromatin accessibility and gene expression data integration via Cycle-consistent Adversarial Network"

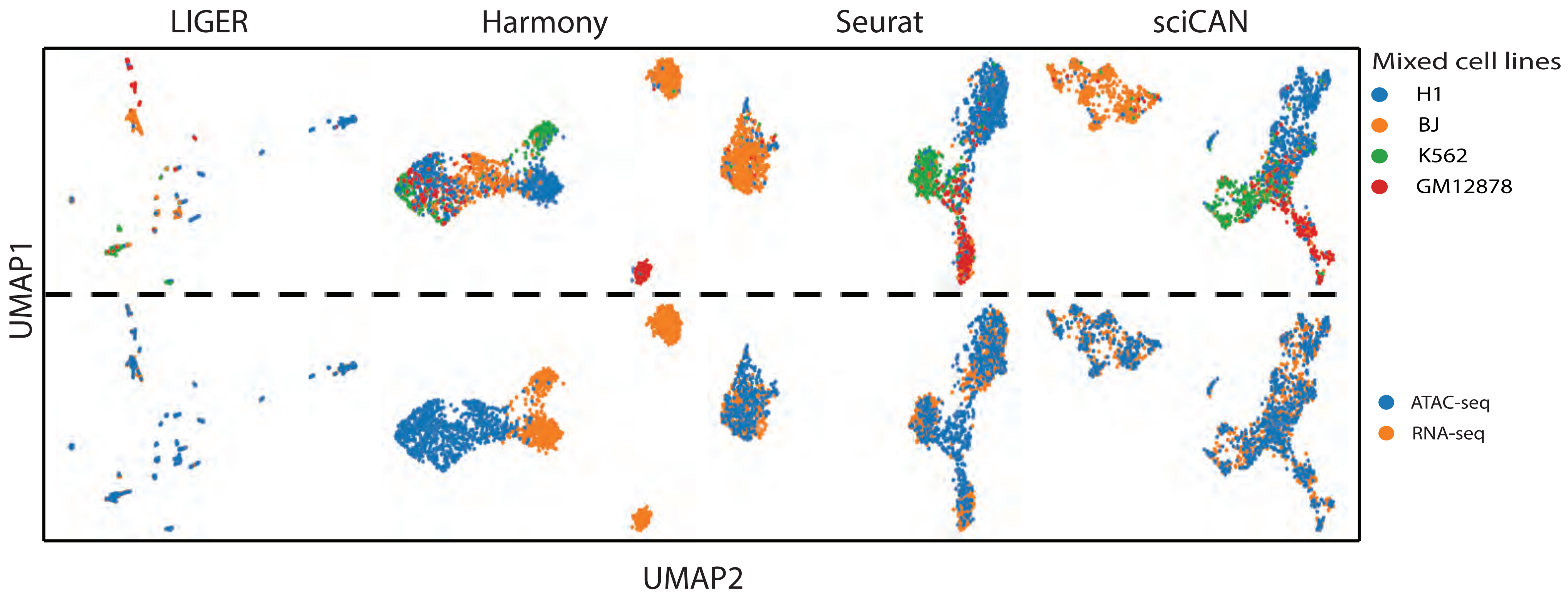

LIGER
Harmony
Seurat
sciCAN
UMAP1
UMAP2
Mixed cell lines
H1
BJ
K562
GM12878
ATAC-seq
RNA-seq

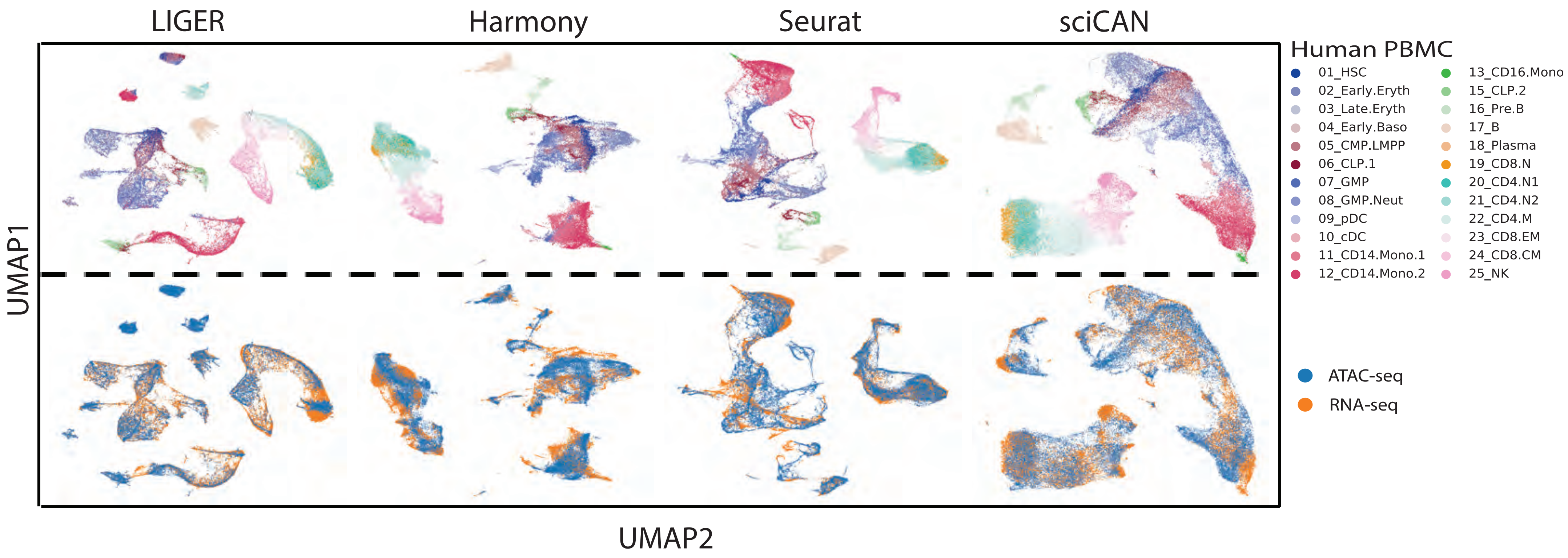
LIGER
Harmony
Seurat
sciCAN
UMAP1
UMAP2
Human PBMC
01_HSC
02_Early.Eryth
03_Late.Eryth
04_Early.Baso
05_CMP.LMPP
06_CLP.1
07_GMP
08_GMP.Neut
09_pDC
10_cDC
11_CD14.Mono.1
12_CD14.Mono.2
13_CD16.Mono
15_CLP.2
16_Pre.B
17_B
18_Plasma
19_CD8.N
20_CD4.N1
21_CD4.N2
22_CD4.M
23_CD8.EM
24_CD8.CM
25_NK
ATAC-seq
RNA-seq

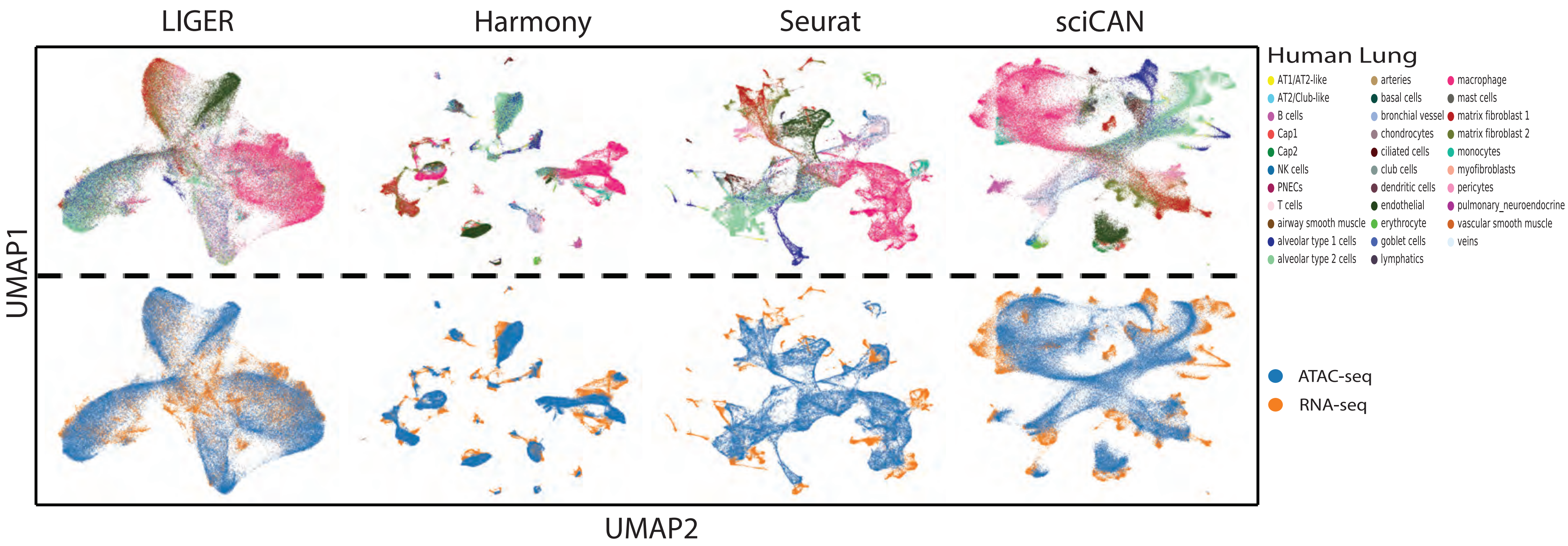
LIGER
Harmony
Seurat
sciCAN
Human Lung
AT1/AT2-like
AT2/Club-like
B cells
Cap1
Cap2
NK cells
PNECs
T cells
airway smooth muscle
alveolar type 1 cells
alveolar type 2 cells
arteries
basal cells
bronchial vessel
chondrocytes
ciliated cells
club cells
dendritic cells
endothelial
erythrocyte
goblet cells
lymphatics
macrophage
mast cells
matrix fibroblast 1
matrix fibroblast 2
monocytes
myofibroblasts
pericytes
pulmonary_neuroendocrine
vascular smooth muscle
veins
ATAC-seq
RNA-seq
UMAP1
UMAP2

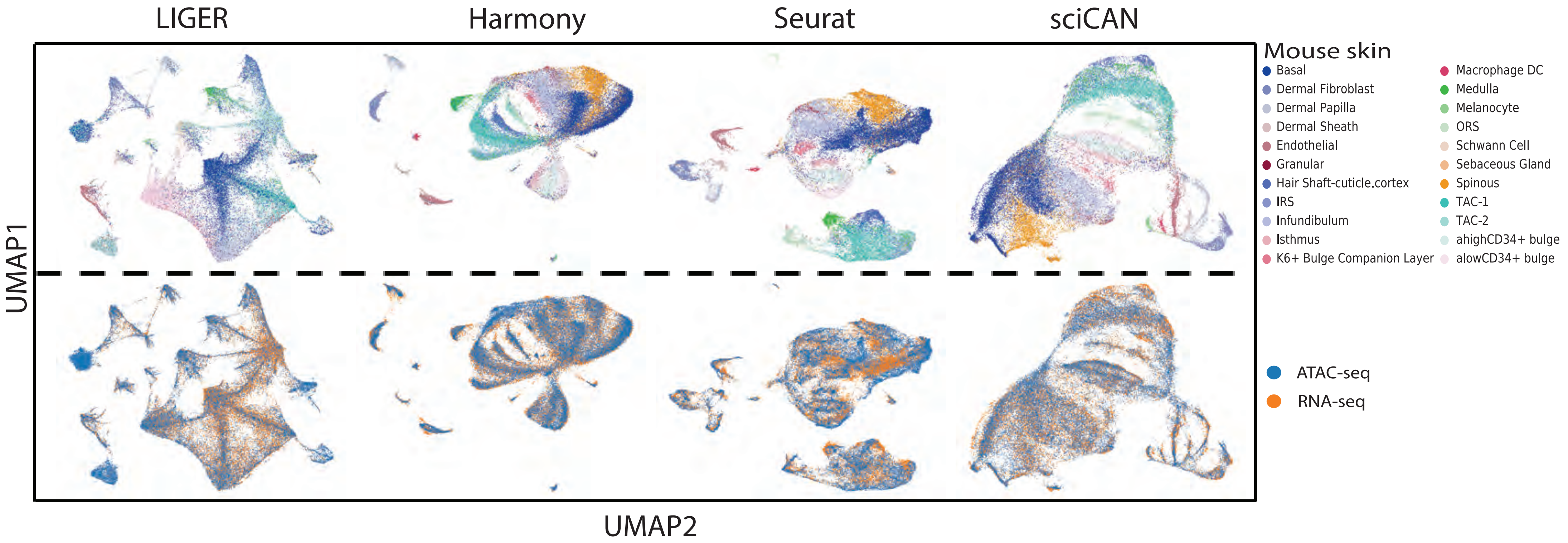
LIGER
Harmony
Seurat
sciCAN
Mouse skin
Basal
Dermal Fibroblast
Dermal Papilla
Dermal Sheath
Endothelial
Granular
Hair Shaft-cuticle.cortex
IRS
Infundibulum
Isthmus
K6+ Bulge Companion Layer
Macrophage DC
Medulla
Melanocyte
ORS
Schwann Cell
Sebaceous Gland
Spinous
TAC-1
TAC-2
ahighCD34+ bulge
alowCD34+ bulge
ATAC-seq
RNA-seq
UMAP1
UMAP2

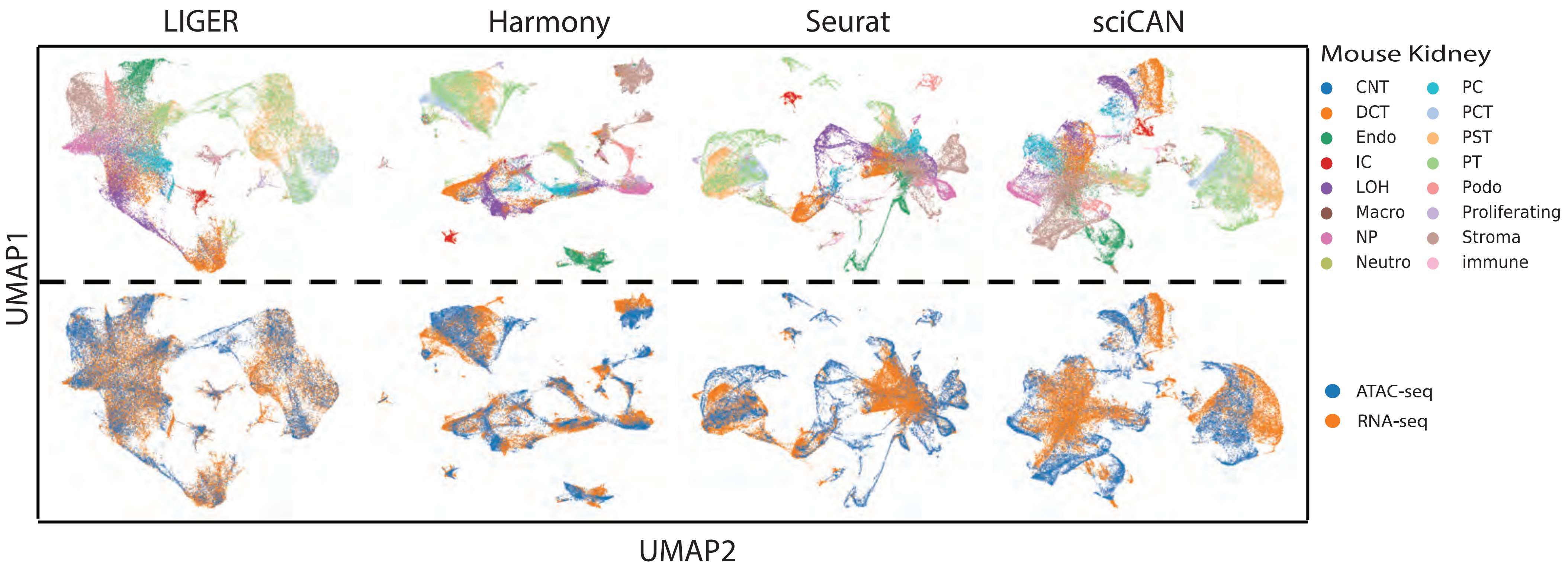

LIGER
Harmony
Seurat
sciCAN
Mouse Kidney
CNT
DCT
Endo
IC
LOH
Macro
NP
Neutro
PC
PCT
PST
PT
Podo
Proliferating
Stroma
immune
ATAC-seq
RNA-seq
UMAP1
UMAP2

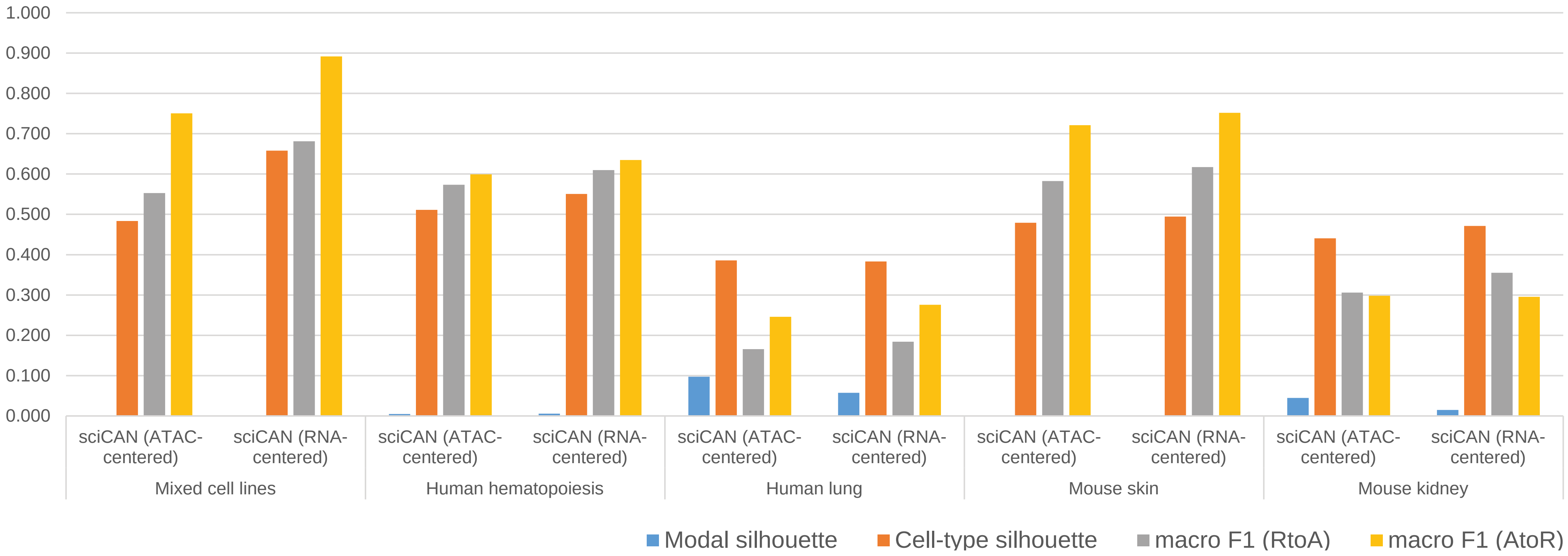

1.000
0.900
0.800
0.700
0.600
0.500
0.400
0.300
0.200
0.100
0.000
sciCAN (ATAC-centered)
sciCAN (RNA-centered)
Mixed cell lines
Human hematopoiesis
Human lung
Mouse skin
Mouse kidney
Modal silhouette
Cell-type silhouette
macro F1 (RtoA)
macro F1 (AtoR)

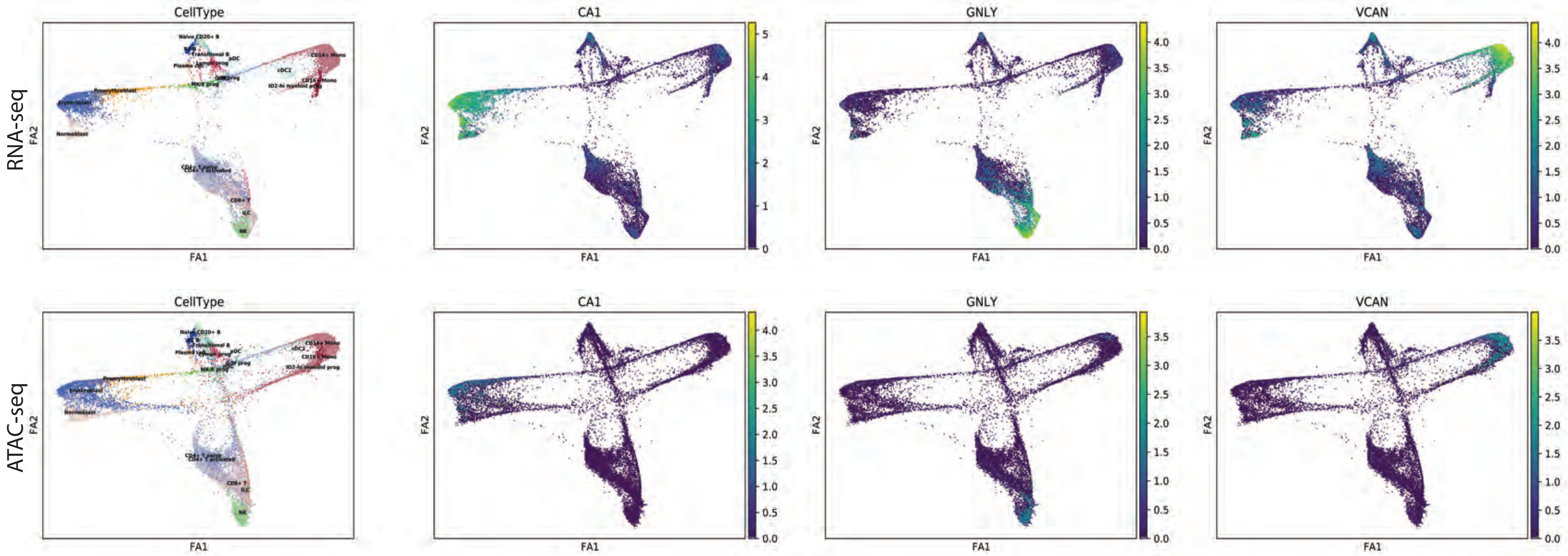
RNA-seq
ATAC-seq
CellType
CA1
GNLY
VCAN
FA1
FA2
Naive CD20+ B
Transitional B
pDC
CD14+ Mono
cDC2
CD16+ Mono
ID2-hi myeloid prog
Proerythroblast
Erythroblast
Normoblast
MK/E prog
CD4+ T naive
CD4+ T activated
CD8+ T
ILC
NK

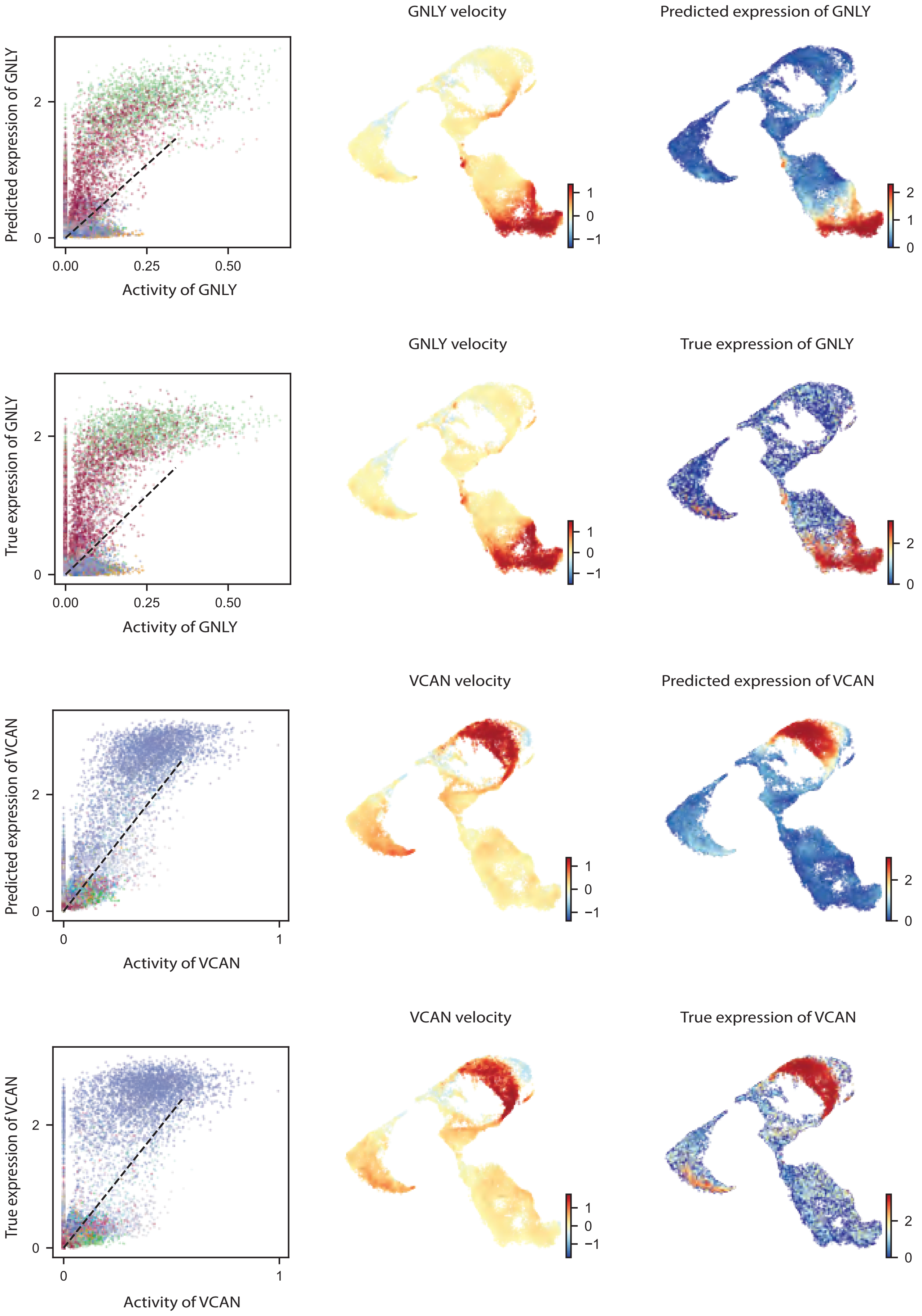
GNLY velocity
Predicted expression of GNLY
Predicted expression of GNLY
Activity of GNLY
0.00
0.25
0.50
0
2
1
0
−1
2
1
0
GNLY velocity
True expression of GNLY
True expression of GNLY
Activity of GNLY
0.00
0.25
0.50
0
2
1
0
−1
2
0
VCAN velocity
Predicted expression of VCAN
Predicted expression of VCAN
Activity of VCAN
0
1
0
2
1
0
−1
2
0
VCAN velocity
True expression of VCAN
True expression of VCAN
Activity of VCAN
0
1
0
2
1
0
−1
2
0

**Supplementary Table 1**

| Data | Sample | # of cells in ATAC-seq | # of cells in RNA-seq | # of cell-types in ATAC-seq | # of cell-types in RNA-seq | Accession | Reference | Processed data |
|---|---|---|---|---|---|---|---|---|
| 1 | Cell lines | 1047 | 1047 | 4 | 4 | GSE126074 | Chen et al., 2019 | |
| 2 | Human hematopoiesis | 33819 | 34901 | 23 | 24 | GSE139369 | Granja et al., 2019 | |
| 3 | Human lung | 82159 | 44294 | 16 | 30 | GSE161383 | Wang et al., 2020 | |
| 4 | Mouse skin | 32231 | 32231 | 22 | 22 | GSE140203 | Ma et al., 2020 | |
| 5 | Mouse kidney | 28316 | 43410 | 11 | 14 | GSE157079 | Miao et al., 2021 | |
| 6 | Human PBMC | 20952 | 20952 | 21 | 21 | NA | NA | https://support.10xgenomics.com/single-cell-multiome-atac-gex/datasets |
| 7 | K562 | 9797 | 22656 | NA | NA | GSE90063/ GSE168851 | Dixit et al., 2016/ Pierce et al., 2021 | |

**Supplementary Table 2**

**RNA-seq**

**C0**

| Rank | Cluster | # cells |
|---|---|---|
| 1 | sgCEP55 | 1727 |
| 2 | sgOGG1 | 1667 |
| 3 | sgPTGER2 | 1637 |
| 4 | sgCABP7 | 1154 |
| 5 | sgCIT | 1141 |
| 6 | sgARHGEF17 | 934 |
| 7 | sgCENPE | 841 |
| 8 | sgAURKC | 836 |
| 9 | sgECT2 | 651 |
| 10 | sgELK1 | 458 |
| 11 | sgAURKB | 380 |
| 12 | sgRACGAP1 | 370 |
| 13 | sgTOR1AIP1 | 281 |
| 14 | sgAURKA | 188 |
| 15 | sgELF1 | 16 |
| 16 | sgEGR1 | 13 |
| 17 | sgE2F4 | 7 |
| 18 | sgNR2C2 | 6 |
| 19 | sgYY1 | 6 |
| 20 | sgGABPA | 5 |
| 21 | sgCREB1 | 4 |
| 22 | sgIRF1 | 4 |
| 23 | sgETS1 | 1 |
| | SUM | 12327 |

**C1**

| Rank | Cluster | # cells |
|---|---|---|
| 1 | sgELK1 | 1615 |
| 2 | sgELF1 | 1296 |
| 3 | sgCREB1 | 882 |
| 4 | sgEGR1 | 787 |
| 5 | sgETS1 | 641 |
| 6 | sgYY1 | 604 |
| 7 | sgGABPA | 573 |
| 8 | sgNR2C2 | 565 |
| 9 | sgE2F4 | 491 |
| 10 | sgIRF1 | 490 |
| 11 | sgPTGER2 | 45 |
| 12 | sgCEP55 | 41 |
| 13 | sgOGG1 | 39 |
| 14 | sgCABP7 | 37 |
| 15 | sgARHGEF17 | 24 |
| 16 | sgCENPE | 18 |
| 17 | sgCIT | 18 |
| 18 | sgAURKC | 15 |
| 19 | sgECT2 | 13 |
| 20 | sgTOR1AIP1 | 12 |
| 21 | sgAURKA | 11 |
| 22 | sgAURKB | 8 |
| 23 | sgRACGAP1 | 8 |
| | SUM | 8233 |

**C2**

| Rank | Cluster | # cells |
|---|---|---|
| 1 | sgCEP55 | 300 |
| 2 | sgPTGER2 | 291 |
| 3 | sgOGG1 | 251 |
| 4 | sgCABP7 | 203 |
| 5 | sgCIT | 201 |
| 6 | sgARHGEF17 | 153 |
| 7 | sgCENPE | 152 |
| 8 | sgAURKC | 149 |
| 9 | sgECT2 | 110 |
| 10 | sgELK1 | 81 |
| 11 | sgRACGAP1 | 68 |
| 12 | sgAURKB | 56 |
| 13 | sgTOR1AIP1 | 51 |
| 14 | sgAURKA | 28 |
| 15 | sgELF1 | 1 |
| 16 | sgNR2C2 | 1 |
| 17 | sgCREB1 | 0 |
| 18 | sgE2F4 | 0 |
| 19 | sgEGR1 | 0 |
| 20 | sgETS1 | 0 |
| 21 | sgGABPA | 0 |
| 22 | sgIRF1 | 0 |
| 23 | sgYY1 | 0 |
| | SUM | 2096 |

**RNA-seq**

**C0**

| Rank | Cluster | Proportion |
|---|---|---|
| 1 | sgCEP55 | 0.1401 |
| 2 | sgOGG1 | 0.1352 |
| 3 | sgPTGER2 | 0.1328 |
| 4 | sgCABP7 | 0.0936 |

**C1**

| Rank | Cluster | Proportion |
|---|---|---|
| 1 | sgELK1 | 0.1962 |
| 2 | sgELF1 | **0.1574** |
| 3 | sgCREB1 | 0.1071 |
| 4 | sgEGR1 | 0.0956 |

**C2**

| Rank | Cluster | Proportion |
|---|---|---|
| 1 | sgCEP55 | 0.1431 |
| 2 | sgPTGER2 | 0.1388 |
| 3 | sgOGG1 | 0.1198 |
| 4 | sgCABP7 | 0.0969 |

| | | |
|---|---|---|
| 5 | sgCIT | 0.0926 |
| 6 | sgARHGEF17 | 0.0758 |
| 7 | sgCENPE | 0.0682 |
| 8 | sgAURKC | 0.0678 |
| 9 | sgECT2 | 0.0528 |
| 10 | sgELK1 | 0.0372 |
| 11 | sgAURKB | 0.0308 |
| 12 | sgRACGAP1 | 0.0300 |
| 13 | sgTOR1AIP1 | 0.0228 |
| 14 | sgAURKA | 0.0153 |
| 15 | sgELF1 | **0.0013** |
| 16 | sgEGR1 | 0.0011 |
| 17 | sgE2F4 | 0.0006 |
| 18 | sgNR2C2 | 0.0005 |
| 19 | sgYY1 | **0.0005** |
| 20 | sgGABPA | **0.0004** |
| 21 | sgCREB1 | 0.0003 |
| 22 | sgIRF1 | 0.0003 |
| 23 | sgETS1 | 0.0001 |

| | | |
|---|---|---|
| 5 | sgETS1 | 0.0779 |
| 6 | sgYY1 | **0.0734** |
| 7 | sgGABPA | **0.0696** |
| 8 | sgNR2C2 | 0.0686 |
| 9 | sgE2F4 | 0.0596 |
| 10 | sgIRF1 | 0.0595 |
| 11 | sgPTGER2 | 0.0055 |
| 12 | sgCEP55 | 0.0050 |
| 13 | sgOGG1 | 0.0047 |
| 14 | sgCABP7 | 0.0045 |
| 15 | sgARHGEF17 | 0.0029 |
| 16 | sgCENPE | 0.0022 |
| 17 | sgCIT | 0.0022 |
| 18 | sgAURKC | 0.0018 |
| 19 | sgECT2 | 0.0016 |
| 20 | sgTOR1AIP1 | 0.0015 |
| 21 | sgAURKA | 0.0013 |
| 22 | sgAURKB | 0.0010 |
| 23 | sgRACGAP1 | 0.0010 |

| | | |
|---|---|---|
| 5 | sgCIT | 0.0959 |
| 6 | sgARHGEF17 | 0.0730 |
| 7 | sgCENPE | 0.0725 |
| 8 | sgAURKC | 0.0711 |
| 9 | sgECT2 | 0.0525 |
| 10 | sgELK1 | 0.0386 |
| 11 | sgRACGAP1 | 0.0324 |
| 12 | sgAURKB | 0.0267 |
| 13 | sgTOR1AIP1 | 0.0243 |
| 14 | sgAURKA | 0.0134 |
| 15 | sgELF1 | **0.0005** |
| 16 | sgNR2C2 | 0.0005 |
| 17 | sgCREB1 | 0.0000 |
| 18 | sgE2F4 | 0.0000 |
| 19 | sgEGR1 | 0.0000 |
| 20 | sgETS1 | 0.0000 |
| 21 | sgGABPA | **0.0000** |
| 22 | sgIRF1 | 0.0000 |
| 23 | sgYY1 | **0.0000** |

**ATAC-seq**

**C0**

| Rank | Cluster | # cells |
|---|---|---|
| 1 | sgFOSL1 | 193 |
| 2 | sgSETDB1 | 187 |
| 3 | sgELF1 | 164 |
| 4 | sgCEBPZ | 163 |
| 5 | sgPBX2 | 162 |
| 6 | sgZBTB11 | 162 |
| 7 | sgATF1 | 161 |
| 8 | sgTRIM28 | 161 |
| 9 | sgCUX1 | 160 |
| 10 | sgBCLAF1 | 159 |
| 11 | sgTBP | 159 |
| 12 | sgCEBPB | 157 |

**C1**

| Rank | Cluster | # cells |
|---|---|---|
| 1 | sgZNF280A | 122 |
| 2 | sgTFDP1 | 113 |
| 3 | sgELF1 | 111 |
| 4 | sgNFYB | 107 |
| 5 | sgATF1 | 103 |
| 6 | sgHINFP | 103 |
| 7 | sgCEBPZ | 100 |
| 8 | sgCUX1 | 100 |
| 9 | sgZNF407 | 99 |
| 10 | sgBCLAF1 | 97 |
| 11 | sgZBTB11 | 96 |
| 12 | sgBRF2 | 94 |

**C2**

| Rank | Cluster | # cells |
|---|---|---|
| 1 | sgTRIM28 | 58 |
| 2 | sgGATA1 | 50 |
| 3 | sgATF1 | 49 |
| 4 | sgZNF280A | 48 |
| 5 | sgZNF407 | 47 |
| 6 | sgPBX2 | 42 |
| 7 | sgCEBPZ | 41 |
| 8 | sgMAX | 41 |
| 9 | sgTFDP1 | 41 |
| 10 | sgFOSL1 | 40 |
| 11 | sgELF1 | 39 |
| 12 | sgGABPA | 39 |

| | | |
|---|---|---|
| 13 | sgBRF2 | 155 |
| 14 | sgHINFP | 152 |
| 15 | sgRPL9 | 152 |
| 16 | sgZNF280A | 152 |
| 17 | sgNFYB | 151 |
| 18 | sgNFE2 | 150 |
| 19 | sgZNF407 | 142 |
| 20 | sgTFDP1 | 140 |
| 21 | sgPOLR1D | 139 |
| 22 | sgGABPA | 136 |
| 23 | sgREST | 134 |
| 24 | sgYY1 | 134 |
| 25 | sgZZZ3 | 133 |
| 26 | sgARID2 | 130 |
| 27 | sgCTCF | 120 |
| 28 | sgTHAP1 | 120 |
| 29 | sgNRF1 | 119 |
| 30 | sgHSPA5 | 109 |
| 31 | sgMAX | 105 |
| 32 | sgKLF16 | 95 |
| 33 | sgATF3 | 92 |
| 34 | sgMYC | 88 |
| 35 | sgARID3A | 87 |
| 36 | sgGTF2B | 76 |
| 37 | sgCDC5L | 75 |
| 38 | sgKLF1 | 66 |
| 39 | sgCAD | 61 |
| 40 | sgGATA1 | 44 |
| | SUM | 5245 |

| | | |
|---|---|---|
| 13 | sgPBX2 | 94 |
| 14 | sgTBP | 93 |
| 15 | sgTRIM28 | 90 |
| 16 | sgREST | 89 |
| 17 | sgCEBPB | 87 |
| 18 | sgSETDB1 | 85 |
| 19 | sgZZZ3 | 82 |
| 20 | sgFOSL1 | 80 |
| 21 | sgNRF1 | 80 |
| 22 | sgARID2 | 78 |
| 23 | sgNFE2 | 74 |
| 24 | sgKLF16 | 72 |
| 25 | sgCAD | 71 |
| 26 | sgCDC5L | 71 |
| 27 | sgMAX | 71 |
| 28 | sgYY1 | 70 |
| 29 | sgCTCF | 69 |
| 30 | sgPOLR1D | 67 |
| 31 | sgHSPA5 | 64 |
| 32 | sgARID3A | 59 |
| 33 | sgKLF1 | 59 |
| 34 | sgGTF2B | 54 |
| 35 | sgGATA1 | 52 |
| 36 | sgMYC | 51 |
| 37 | sgGABPA | 49 |
| 38 | sgTHAP1 | 49 |
| 39 | sgRPL9 | 39 |
| 40 | sgATF3 | 37 |
| | SUM | 3181 |

| | | |
|---|---|---|
| 13 | sgSETDB1 | 39 |
| 14 | sgTBP | 39 |
| 15 | sgZZZ3 | 39 |
| 16 | sgNRF1 | 38 |
| 17 | sgZBTB11 | 38 |
| 18 | sgCUX1 | 37 |
| 19 | sgBRF2 | 36 |
| 20 | sgCEBPB | 34 |
| 21 | sgHSPA5 | 34 |
| 22 | sgKLF1 | 34 |
| 23 | sgNFE2 | 34 |
| 24 | sgNFYB | 34 |
| 25 | sgREST | 33 |
| 26 | sgPOLR1D | 31 |
| 27 | sgARID2 | 30 |
| 28 | sgBCLAF1 | 28 |
| 29 | sgCDC5L | 28 |
| 30 | sgTHAP1 | 28 |
| 31 | sgHINFP | 27 |
| 32 | sgCTCF | 24 |
| 33 | sgYY1 | 24 |
| 34 | sgATF3 | 23 |
| 35 | sgGTF2B | 22 |
| 36 | sgKLF16 | 22 |
| 37 | sgRPL9 | 22 |
| 38 | sgCAD | 21 |
| 39 | sgARID3A | 20 |
| 40 | sgMYC | 17 |
| | SUM | 1371 |

**ATAC-seq**

**C0**

| Rank | Cluster | Proportion |
|---|---|---|
| 1 | sgFOSL1 | 0.0368 |
| 2 | sgSETDB1 | 0.0357 |
| 3 | sgELF1 | **0.0313** |

**C1**

| Rank | Cluster | Proportion |
|---|---|---|
| 1 | sgZNF280A | 0.0384 |
| 2 | sgTFDP1 | 0.0355 |
| 3 | sgELF1 | **0.0349** |

**C2**

| Rank | Cluster | Proportion |
|---|---|---|
| 1 | sgTRIM28 | 0.0423 |
| 2 | sgGATA1 | 0.0365 |
| 3 | sgATF1 | 0.0357 |

| | | |
|---|---|---|
| 4 | sgCEBPZ | 0.0311 |
| 5 | sgPBX2 | 0.0309 |
| 6 | sgZBTB11 | 0.0309 |
| 7 | sgATF1 | 0.0307 |
| 8 | sgTRIM28 | 0.0307 |
| 9 | sgCUX1 | 0.0305 |
| 10 | sgBCLAF1 | 0.0303 |
| 11 | sgTBP | 0.0303 |
| 12 | sgCEBPB | 0.0299 |
| 13 | sgBRF2 | 0.0296 |
| 14 | sgHINFP | 0.0290 |
| 15 | sgRPL9 | 0.0290 |
| 16 | sgZNF280A | 0.0290 |
| 17 | sgNFYB | 0.0288 |
| 18 | sgNFE2 | 0.0286 |
| 19 | sgZNF407 | 0.0271 |
| 20 | sgTFDP1 | 0.0267 |
| 21 | sgPOLR1D | 0.0265 |
| 22 | sgGABPA | **0.0259** |
| 23 | sgREST | 0.0255 |
| 24 | sgYY1 | **0.0255** |
| 25 | sgZZZ3 | 0.0254 |
| 26 | sgARID2 | 0.0248 |
| 27 | sgCTCF | 0.0229 |
| 28 | sgTHAP1 | 0.0229 |
| 29 | sgNRF1 | 0.0227 |
| 30 | sgHSPA5 | 0.0208 |
| 31 | sgMAX | 0.0200 |
| 32 | sgKLF16 | 0.0181 |
| 33 | sgATF3 | 0.0175 |
| 34 | sgMYC | 0.0168 |
| 35 | sgARID3A | 0.0166 |
| 36 | sgGTF2B | 0.0145 |
| 37 | sgCDC5L | 0.0143 |
| 38 | sgKLF1 | 0.0126 |

| | | |
|---|---|---|
| 4 | sgNFYB | 0.0336 |
| 5 | sgATF1 | 0.0324 |
| 6 | sgHINFP | 0.0324 |
| 7 | sgCEBPZ | 0.0314 |
| 8 | sgCUX1 | 0.0314 |
| 9 | sgZNF407 | 0.0311 |
| 10 | sgBCLAF1 | 0.0305 |
| 11 | sgZBTB11 | 0.0302 |
| 12 | sgBRF2 | 0.0296 |
| 13 | sgPBX2 | 0.0296 |
| 14 | sgTBP | 0.0292 |
| 15 | sgTRIM28 | 0.0283 |
| 16 | sgREST | 0.0280 |
| 17 | sgCEBPB | 0.0273 |
| 18 | sgSETDB1 | 0.0267 |
| 19 | sgZZZ3 | 0.0258 |
| 20 | sgFOSL1 | 0.0251 |
| 21 | sgNRF1 | 0.0251 |
| 22 | sgARID2 | 0.0245 |
| 23 | sgNFE2 | 0.0233 |
| 24 | sgKLF16 | 0.0226 |
| 25 | sgCAD | 0.0223 |
| 26 | sgCDC5L | 0.0223 |
| 27 | sgMAX | 0.0223 |
| 28 | sgYY1 | **0.0220** |
| 29 | sgCTCF | 0.0217 |
| 30 | sgPOLR1D | 0.0211 |
| 31 | sgHSPA5 | 0.0201 |
| 32 | sgARID3A | 0.0185 |
| 33 | sgKLF1 | 0.0185 |
| 34 | sgGTF2B | 0.0170 |
| 35 | sgGATA1 | 0.0163 |
| 36 | sgMYC | 0.0160 |
| 37 | sgGABPA | **0.0154** |
| 38 | sgTHAP1 | 0.0154 |

| | | |
|---|---|---|
| 4 | sgZNF280A | 0.0350 |
| 5 | sgZNF407 | 0.0343 |
| 6 | sgPBX2 | 0.0306 |
| 7 | sgCEBPZ | 0.0299 |
| 8 | sgMAX | 0.0299 |
| 9 | sgTFDP1 | 0.0299 |
| 10 | sgFOSL1 | 0.0292 |
| 11 | sgELF1 | **0.0284** |
| 12 | sgGABPA | **0.0284** |
| 13 | sgSETDB1 | 0.0284 |
| 14 | sgTBP | 0.0284 |
| 15 | sgZZZ3 | 0.0284 |
| 16 | sgNRF1 | 0.0277 |
| 17 | sgZBTB11 | 0.0277 |
| 18 | sgCUX1 | 0.0270 |
| 19 | sgBRF2 | 0.0263 |
| 20 | sgCEBPB | 0.0248 |
| 21 | sgHSPA5 | 0.0248 |
| 22 | sgKLF1 | 0.0248 |
| 23 | sgNFE2 | 0.0248 |
| 24 | sgNFYB | 0.0248 |
| 25 | sgREST | 0.0241 |
| 26 | sgPOLR1D | 0.0226 |
| 27 | sgARID2 | 0.0219 |
| 28 | sgBCLAF1 | 0.0204 |
| 29 | sgCDC5L | 0.0204 |
| 30 | sgTHAP1 | 0.0204 |
| 31 | sgHINFP | 0.0197 |
| 32 | sgCTCF | 0.0175 |
| 33 | sgYY1 | **0.0175** |
| 34 | sgATF3 | 0.0168 |
| 35 | sgGTF2B | 0.0160 |
| 36 | sgKLF16 | 0.0160 |
| 37 | sgRPL9 | 0.0160 |
| 38 | sgCAD | 0.0153 |

| | | |
|---|---|---|
| 39 | sgCAD | 0.0116 |
| 40 | sgGATA1 | 0.0084 |

| | | |
|---|---|---|
| 39 | sgRPL9 | 0.0123 |
| 40 | sgATF3 | 0.0116 |

| | | |
|---|---|---|
| 39 | sgARID3A | 0.0146 |
| 40 | sgMYC | 0.0124 |
